# Extracellular Ca^2+^-sensitive fluorescent protein biosensor based on a collagen-binding domain

**DOI:** 10.1101/2020.03.11.987446

**Authors:** Irina A. Okkelman, Ryan McGarrigle, Shane O’Carroll, Daniel Carvajal Berrio, Katja Schenke-Layland, James Hynes, Ruslan I. Dmitriev

**Affiliations:** Metabolic Imaging Group, Laboratory of Biophysics and Bioanalysis, ABCRF, University College Cork, College Road, Cork, Ireland; Agilent Technologies Ireland Limited, Little Island, Cork; Department of Women’s Health, Research Institute for Women’s Health, Eberhard Karls University Tübingen, Tübingen, Germany; Cluster of Excellence iFIT (EXC 2180) “Image-Guided and Functionally Instructed Tumor; NMI Natural and Medical Sciences Institute at the University of Tübingen, Reutlingen, Germany; Department of Medicine/Cardiology, Cardiovascular Research Laboratories, David Geffen School of Medicine at UCLA, Los Angeles, CA, USA; I.M. Sechenov First Moscow State University, Institute for Regenerative Medicine, Moscow, Russian Federation

## Abstract

The importance of extracellular gradients of biomolecules becomes increasingly appreciated in the processes of tissue development and regeneration, in health and disease. In particular, dynamics of extracellular calcium concentration is rarely studied. Here, we present low affinity Ca^2+^ biosensor based on Twitch-2B fluorescent protein fused with the cellulose- and collagen-binding peptides. These recombinant chimeric proteins can bind cellulose and collagen scaffolds and enable for scaffold-based biosensing of Ca^2+^ in proximity of live 3D tissue models. We found that the Twitch-2B mutant is compatible with intensity-based ratiometric and fluorescence lifetime imaging microscopy (FLIM) measurement formats, under one- and two-photon excitation modes. Furthermore, the donor fluorescence lifetime of ColBD-Twitch displays response to [Ca^2+^] over a range of ∼2-2.5 ns, making it attractive biosensor for multiplexed FLIM microscopy assays. To evaluate performance of this biosensor in physiological measurements, we applied ColBD-Twitch to the live Lgr5-GFP mouse intestinal organoid culture and measured its responses to the changes in extracellular Ca^2+^ upon chelation with EGTA. When we combined it with spectrally resolved FLIM of lipid droplets using Nile Red dye, we observed changes in cytoplasmic and basal membrane-associated lipid droplet composition in response to the extracellular Ca^2+^ depletion, suggesting that intestinal epithelium can respond to and compensate such treatment. Altogether, our results demonstrate ColBD-Twitch as a prospective Ca^2+^ sensor for multiplexed FLIM analysis in a complex 3D tissue environment.

## Introduction

Extracellular cues become increasingly important to understand tissue organization and development. For instance, building-up the tissue from the stem cell-derived organoids demands a deep understanding of the time-dependent changes of biophysical parameters experienced during growth and differentiation [1-3]. In addition to the physical tension [4] and the extracellular matrix itself [5], changing cell density, composition and organization can also affect gradients of O_2_ (hypoxia) [6, 7], distribution and diffusion of reactive oxygen species, peptides and growth factors, pH and other ions such as Ca^2+^ [8, 9].

A number of reports have indicated the role of extracellular Ca^2+^ as one the primary signals that influence cell function [10]: thus, regulation of extracellular Ca^2+^ have been shown to be relevant to gastrointestinal tract function [11, 12], bone marrow [13], brain and CNS function [14, 15], smooth and skeletal muscles [16], wound healing [17], plasma membrane repair [18], engineered organoid-like tissues [19] and microbial biofilm formation [20]. Every mammalian cell relies on the Ca^2+^ homeostasis, which in turn is tightly regulated through the activities of channels, ATPases, Na^+^/ Ca^2+^ exchanger, cytosolic Ca^2+^-binding proteins and various depots such as endoplasmic reticulum, secretory vesicles and mitochondria [21-24]. Calcium influx and efflux processes through the plasma membrane are tightly regulated and the ability to sense peri- / extracellular Ca^2+^ dynamics can become a valuable tool for probing cell function without the need of labeling it.

However, Ca^2+^ sensors with affinity in range of 0-10 mM are rarely realized in optical imaging and analytical measurements [25-28]. The main challenges in sensing extracellular Ca^2+^ ([Ca^2+^]_o_) with most of experimental *in vitro* cell-based models are: (i) relatively large extracellular space volume surrounding the cells, resulting in quick equilibrium of Ca^2+^ across the whole volume; (ii) lack of suitable fluorescent probes and protein biosensors [10, 29].

Biosensor scaffold materials based on the labeling of solid and gel-based materials, can be employed in cell and tissue engineering and provide additional chemo-/ biosensing functionalities [7, 30-41]. The sensing molecule is deposited at or attached to the fiber or scaffold material and not diffusely distributed across the sample; in some cases this approach is very similar to the use of O_2_-sensing films and foils [42-45]. Materials employed in production of scaffolds for the tissue engineering can be either synthetic (e.g. polymer-based) and bio-/ macromolecule-based, e.g. produced from natural hydrogels, decellularized materials and other moieties. Thus, different approaches are needed for the deposition of sensing molecules and these can be achieved via physical impregnation / entrapment, covalent conjugation or through the specific biomolecular interactions. Physical entrapment is highly useful with hydrophobic (and sometimes hydrophilic ‘leaky’ dendrimer) dyes and can be provided by electrospinning and solvent-swelling methods [7, 32, 34]. The covalent immobilization was reported for many organic polymers including fluorescent pH-sensing proteins [46, 47]. Recently, specific biorecognition-based binding was described for producing biosensor cellulose materials, with the help of cellulose-binding domain [33]. In this approach, biocompatible scaffold and biosensor protein are used, making it attractive option for the biosensor design.

A great number of FRET-based protein biosensors such as GECI having various structures and origin enable sensing of Ca^2+^ [28, 48-51]. Traditionally, sensing of Ca^2+^ was focused on the temporal resolution of Ca^2+^ fluxes and not on measuring of their magnitude. Troponin-based FRET Ca^2+^-sensing biosensors such as Twitch-2B are highly attractive for quantitative ratiometric and potentially fluorescence lifetime imaging microscopy (FLIM)-based measurements [52-54]. The latter approach, often relying on measurement of fluorescence lifetime or FRET efficiency of the donor fluorescent protein, is promising for quantitative *in vitro* and *in vivo* measurements: it is less susceptible to the differences in tissue light absorption and scattering (which can represent a problem with detecting two different emission wavelengths), often enables absolute calibration and is compatible with two-photon excited deep tissue imaging [52, 54-56].

Recent study reported a number of ‘Twitch’ biosensors with tunable affinity and improved performance in ratiometric detection, with one mutant (Twitch-2B 54S+) showing promising analytical properties for extracellular measurements [53]. Here, we further evaluated its applicability for production of cellulose- and collagen-based extracellular Ca^2+^-sensing scaffold materials and found that it has attractive features for biosensing in both ratiometric and FLIM detection modalities and is useful for multi-parameter imaging of 3D tissue models such as stem cell-derived intestinal organoids.

## Results

### Cloning and production of CBD-Twitch and ColBD-Twitch proteins

Our preliminary research has highlighted the potential for design of scaffold-binding recombinant proteins consisting of cellulose-binding domain with pH (ECFP) or Ca^2+^-sensing (GCaMP2) proteins [33]. Here, we looked at Twitch-2B 54S+ mutant [53] further referred simply as ‘Twitch’, which is based on Cerulean and cpVenus proteins, minimal Ca^2+^-binding domain, and displayed low affinity promising for extracellular measurements. We produced chimeras of Twitch with cellulose- and collagen-binding domains [57, 58] in bacteria and found that CBD-Twitch showed folding and maturation rates resembling to the CBD-ECFP (yields of ∼ 7 mg / L of culture when expressed at 4 °C). In contrast, ColBD-Twitch protein having short von Willebrand Factor (vWF) collagen-binding peptide demonstrated higher yields of the folded protein, achieving > 50-70% folding rates and yield of > 60 mg / L of culture. SDS-PAGE confirmed integrity and purity of the produced proteins although CBD-Twitch displayed presence of the second electrophoretic band (Fig. 1B), not affected by expression conditions in the presence of protease inhibitors (not shown). Purified CBD-Twitch and ColBD-Twitch showed very similar Ca^2+^-sensitivity in solution with nearly twofold ratiometric response to Ca^2+^ (Fig. 1C, D, 2A).

**Figure 1.**
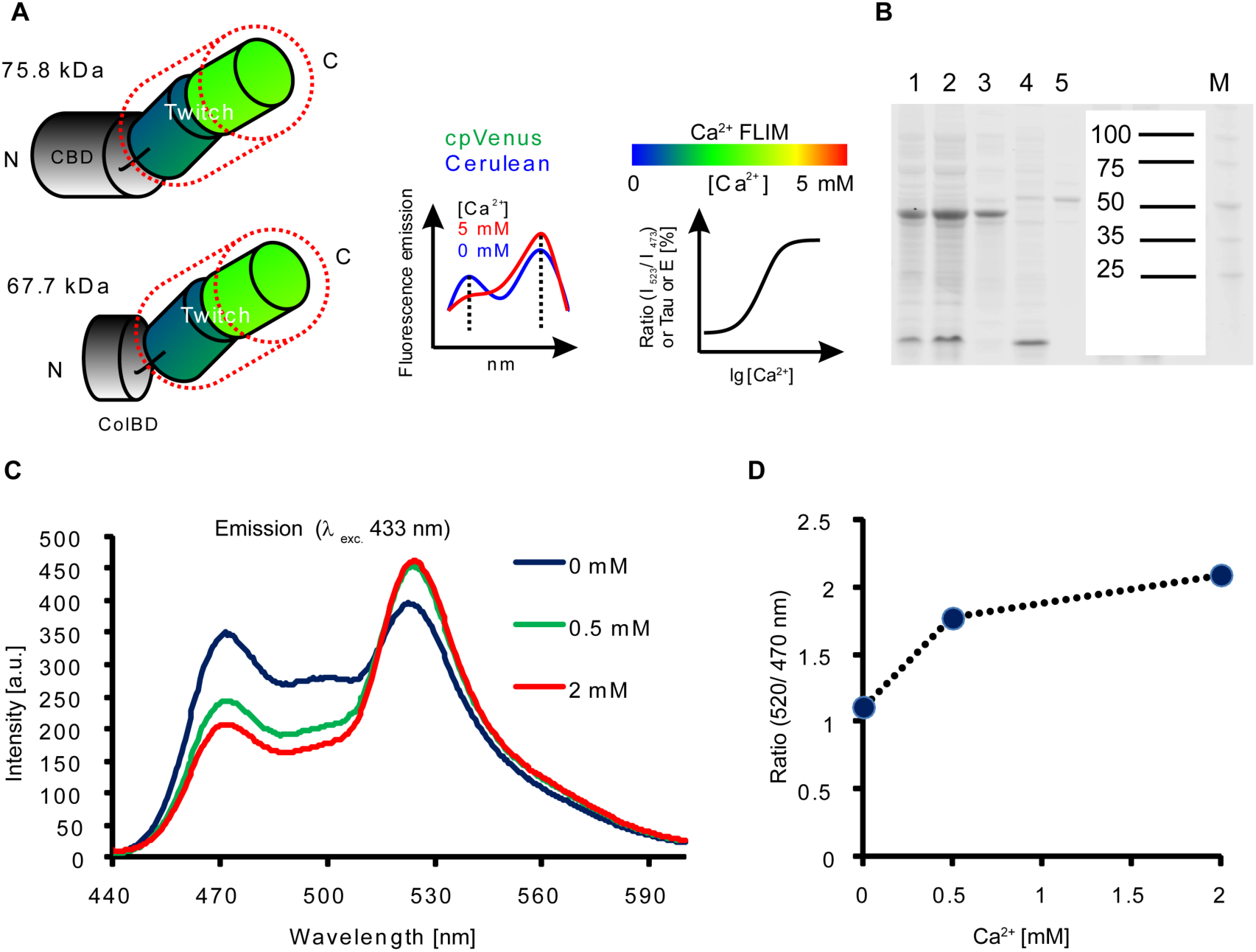
Design and initial evaluation of the cellulose- and collagen-binding domain-containing Twitch-2B 54S+ proteins. A: Schematic structures of recombinant Twitch proteins. Positions of N and C-termini are indicated. Note that N-termini also contain 6xHis sequences. B: SDS-PAGE of crude extracts (1,2,4) and purified ColBD-Twitch (3) and CBD-Twitch (5) proteins. Mw standards are shown on the right. C, D: Fluorescence emission spectra and ratiometric response (ratio of intensities at 520 and 470 nm) for purified CBD-Twitch protein (0.7 μM, 20 °C).

**Figure 2.**
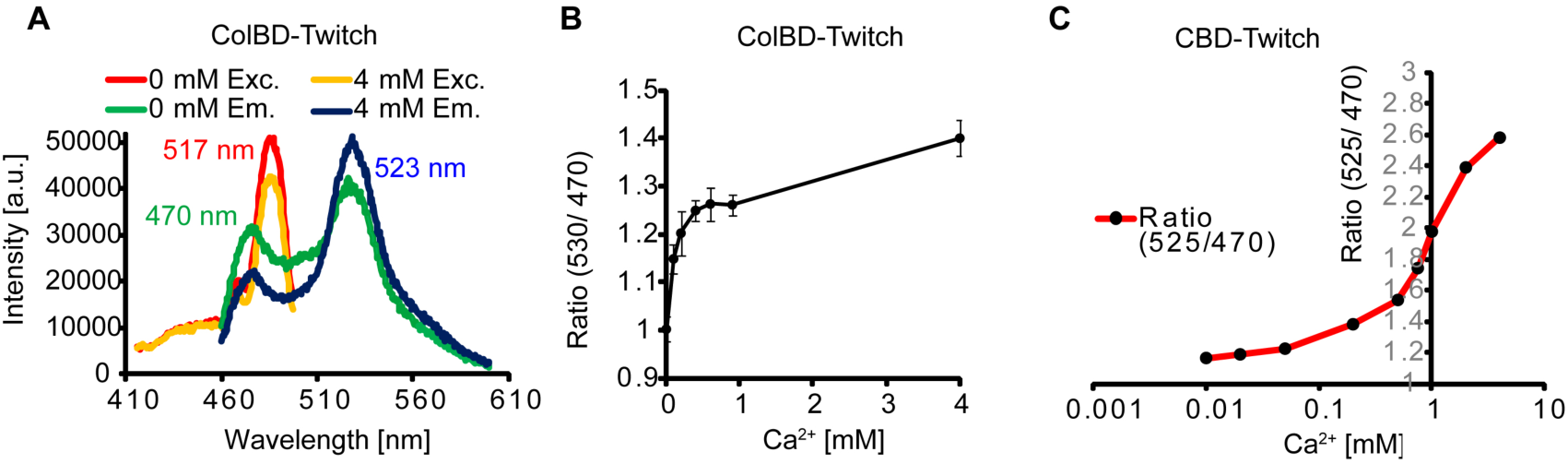
Evaluation of Ca^2+^-dependent response for Twitch proteins in ratiometric detection mode. A: Excitation and emission spectra of ColBD-Twitch at 0 and 4 mM CaCl_2_ in solution, obtained on a microplate reader. Numbers indicate excitation and emission maxima for respective spectra. B: Ratiometric response (emissions at 530 and 470 nm) of ColBD-Twitch across the range of Ca^2+^ concentrations. N=3. C: Ratiometric response to Ca^2+^ for CBD-Twitch in solution, presented in logarithmic scale.

Both proteins displayed binding activity towards cellulose (nanofibrillar and decellularized scaffolds) and collagen (Matrigel, collagen I, IV and decellularized pericardial tissue) correspondingly. For example, ColBD-Twitch rapidly bound collagen I-coated glass microplates, requiring only 15-20 min of the incubation time (Fig. S1).

### Biosensor properties of Twitch proteins in a ratiometric mode

Next we studied biosensing properties of Twitch proteins in a ratiometric intensity-based detection. Using microplate reader with monochromator-based spectral scanning capability, we confirmed ratiometric response of ColBD-Twitch (Fig. 2A). Both CBD- and ColBD-Twitch in solution demonstrated similar [Ca^2+^]-dependent ratiometric calibrations, matching previously reported K_d_ 174 μM and showing approx. 1.4-2fold response between Ca^2+^-free and Ca^2+^-bound forms (Fig. 2B-C).

Given such attractive properties, we looked at the stability of the produced material and its biosensing properties. Thus, we analyzed response to Ca^2+^ for the protein sample stored over 14 days at 4°C (Fig. S2). We detected decreased stability of green channel fluorescence (cpVenus protein) and change of ratio, resulting to decreased performance for the stored protein.

Since the pH-sensitivity of fluorescent proteins is a well-known issue [59], we investigated effects of pH on the Ca^2+^ -sensitivity of the ColBD-Twitch protein (Fig. S3). Comparison of fluorescence intensity for blue (Cerulean) and green (cpVenus) spectral channels at three different concentrations of Ca^2+^ showed that low pH (pH 6.4) significantly affected fluorescence in green channel, thus influencing the ratiometric response. While the range of pH 7-7.4 showed minimal influence of Ca^2+^ -sensitivity, our data indicate that lower pH can indeed affect the performance of this biosensor in the intensity-based detection.

### Evaluation of biosensing properties of Twitch proteins in one- and two-photon excited FLIM

The design of Twitch biosensor implies Ca^2+^ -FRET interaction between Cerulean (donor, blue fluorescence) and cpVenus (acceptor, green fluorescence) proteins, both of which can potentially display changes of the fluorescence lifetime. To understand this phenomenon, we stained Matrigel matrix with ColBD-Twitch and measured the fluorescence lifetime for donor channel using confocal FLIM (Fig. 3). ColBD-Twitch donor fluorescence displayed double exponential decay and decrease of τ_m_ from ∼2.4 to 1.6 ns upon binding Ca^2+^. Changes of fluorescence lifetime and its multi-exponential nature were also confirmed by Phasor analysis (Fig. 3C). We next looked at the broader range of Ca^2+^ concentrations (Fig. 3D, distribution histograms) and observed sensitivity similar to observed in the ratiometric mode (Fig. 2). We also compared according changes of fluorescence lifetime (τ_m_) to the contribution of fraction A1 [%] and FRET efficiency (Fig. 3E,F,G). These results confirmed increase of energy transfer from the donor upon binding Ca^2+^. These data are comparable to the previously reported Cerulean-based Ca^2+^-sensors and suggest that ColBD-Twitch bound to collagen-containing matrix can be used as a Ca^2+^ biosensor.

**Figure 3.**
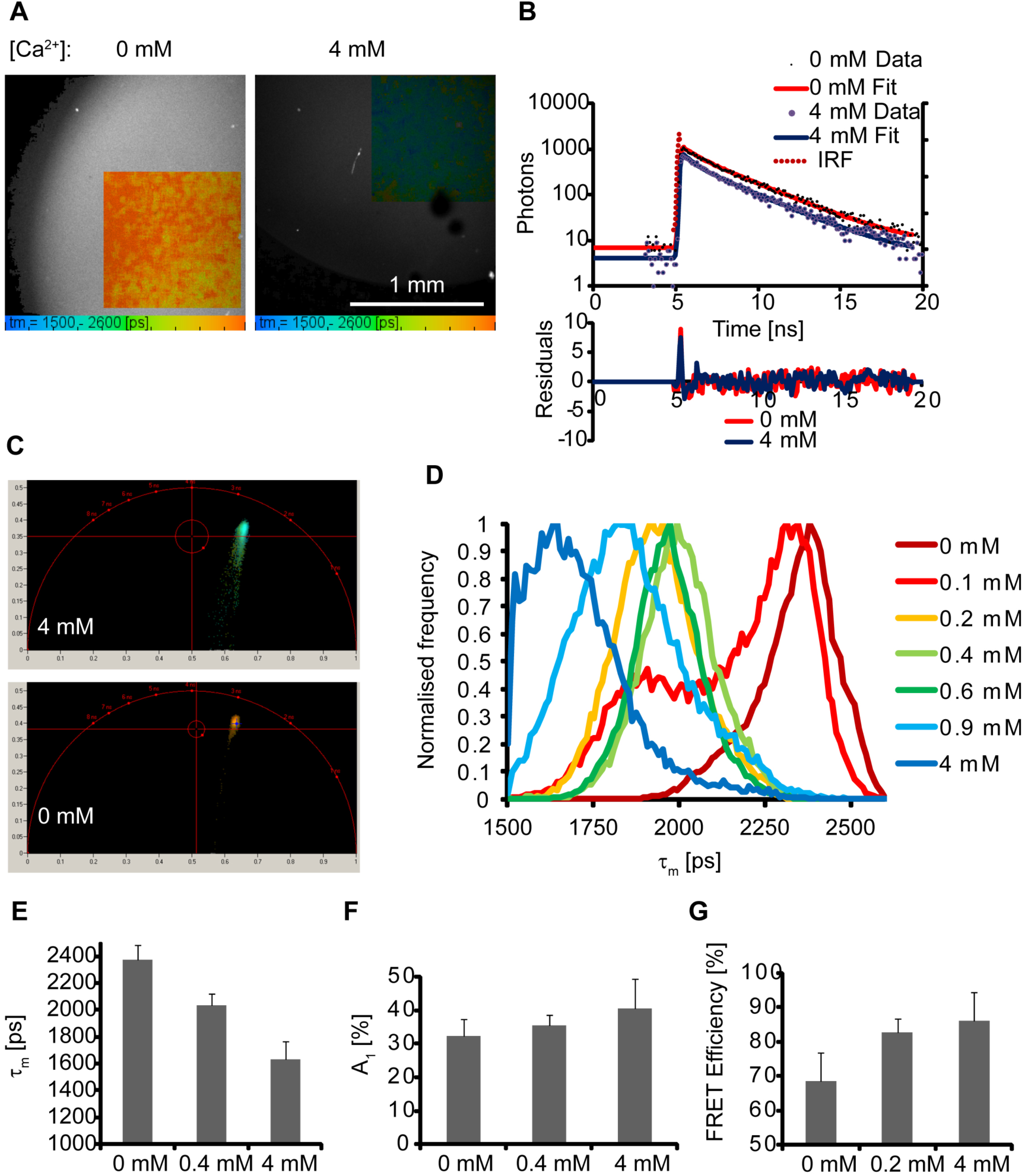
Evaluation of ColBD-Twitch protein as FLIM sensor on a one-photon excited microscope. ColBD-Twitch-stained (1 μM, 3 h) Matrigel matrix was used for measurements in the presence of various concentrations of CaCl_2_ (37 °C). A: Examples of FLIM images for donor channel (405 nm exc., 468 nm em., double-exponential fit, τ_m_) of Matrigel produced at different Ca^2+^ concentrations. B: Example of double-exponential decay fitting. C: Examples of Phasor diagrams. D: Normalized fluorescence lifetime frequency distribution histograms for FLIM images produced at 0-4 mM CaCl_2_. E, F, G: comparisons of τ_m_ (double-exponential fit), A_1_ fraction (%) and FRET efficiency [%] in response to Ca^2+^ binding.

Next, we evaluated Ca^2+^-sensitivity of ColBD-Twitch protein under two-photon excited FLIM, a useful method for *in vivo* applications (Fig. 4). Analysis of two-photon excitation and emission spectra demonstrated difference in Ca^2+^ - dependent responses for donor and acceptor proteins within a Twitch biosensor, with optimal excitation in range of 860-890 nm for both of them. As expected, we observed Ca^2+^-dependent changes in ratiometric intensity mode (Fig. 4A,B), confirming solution-based measurements (Figs. 1-2). In FLIM mode, we saw changes in τ_m_ (donor channel) and observed lifetime changes comparable to one-photon data (Fig. 3). Interestingly, acceptor fluorescence did not reveal drastic changes upon Ca^2+^ binding. We also observed comparable changes in A1 [%] fraction and FRET efficiency in two-photon mode. Thus, ColBD-Twitch protein showed well-resolved changes of fluorescence lifetime of the FRET donor part (blue emitting Cerulean) to changing Ca^2+^ concentrations over the range of 0-4 mM. The double-exponential nature of the fluorescence decay obeyed expected FRET donor mechanics leading to decrease of fluorescence lifetime (due to transfer of energy to the acceptor) upon binding of Ca^2+^, bringing Cerulean and cpVenus to the FRET distance.

**Figure 4.**
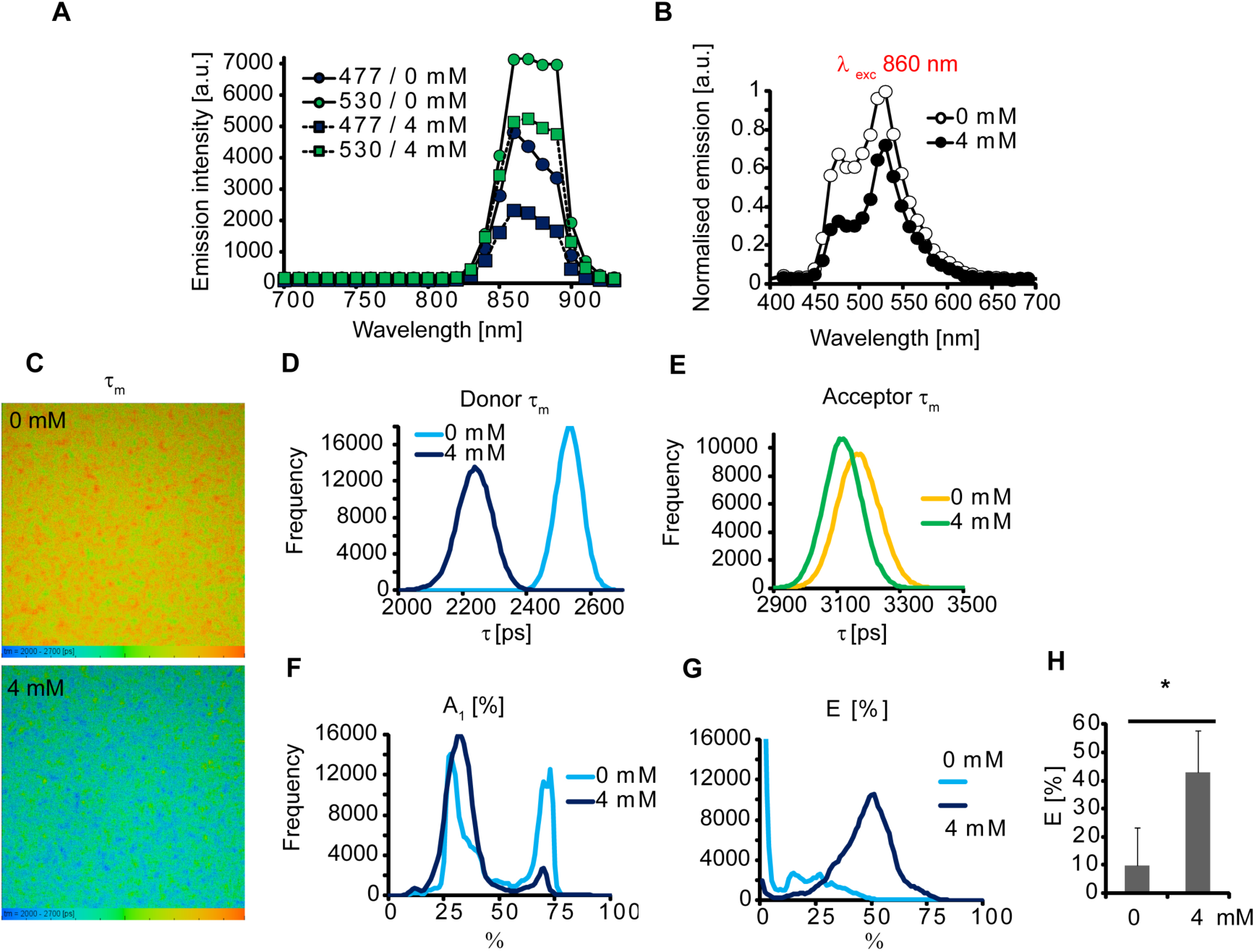
Evaluation of ColBD-Twitch protein in two-photon FLIM. ColBD-Twitch was measured at 0 and 4 mM CaCl_2_ bound to Matrigel (1 μM, 1 h). A, B: two-photon excitation (A) and emission spectra (B). C: FLIM images (double-exponential fit, τ_m_). D, E: lifetime distribution histograms for donor (477 nm em., D) and acceptor (530 nm em., E) spectral channels. F, G, H: FRET efficiency in response to Ca^2+^ expressed in A_1_ [%], F, and FRET ratio (G, H). N=3.

To further understand the behavior of Twitch biosensor with the other types of scaffolds, we performed additional confocal FLIM experiments. First, we produced previously described decellularized (DC) plant tissue celery scaffold [33] and labeled it with CBD-Twitch protein. After observing specific binding, we incubated scaffold with different Ca^2+^ concentrations and using FLIM microscope saw changes in τ_m_ very similar to those obtained with ColBD-Twitch bound to Matrigel, having ∼ 2 to 2.6 ns range of lifetimes for 4 mM to 0.05 mM, respectively (Fig. S4).

We also tested if the Matrigel was not the only one suitable matrix for ColBD-Twitch protein: to do this, we tested if it can bind collagen IV-coated DC celery scaffold (Fig. S5). We found that scaffold displayed slightly different staining pattern than observed with CBD-Twitch on uncoated (pure cellulose) matrix, showing fluorescent signals from cavities rather than from the scaffold. This could be a result of collagen coating of such scaffold and difference in binding between cellulose- and collagen-binding domain proteins. Still, when we analyzed these regions stained with ColBD-Twitch, was also saw expected dynamics of fluorescence lifetime changes. This confirms that ColBD-Twitch can be used with collagen-based scaffolds other than Matrigel (Fig. S5). Collectively, Twitch biosensor fused either with CBD or ColBD domains could bind respective cellulose or collagen-based scaffolds and displayed biosensing properties suitable for one- and two-photon excited FLIM measurements.

### Evaluation of ColBD-Twitch protein in multi-parameter FLIM measurements with the culture of mouse intestinal organoids

Observed spectral and photophysical properties of Twitch biosensor prompted us to test it in a spectrally- and lifetime-resolved FLIM measurements with relevant 3D cell model. We chose mouse Lgr5-GFP small intestinal organoid model [60], which enables live tracing of stem (Lgr5-GFP^+^) and differentiated enterocyte cells, grows in Matrigel matrix and recapitulates most of the functions of the native epithelium [61, 62]. Mouse intestinal organoids have been already extensively evaluated in a number of FLIM and PLIM assays, informing on cell oxygenation, proliferation, mitochondrial polarization, redox status and others [33, 63-67]. Although in previous work we have demonstrated pH-FLIM imaging of organoids combined with O_2_ imaging, we were not able to measure acidification directly in Matrigel and not assessed extracellular Ca^2+^ [33].

First, we tested if we can combine Matrigel-based Ca^2+^ sensing via ColBD-Twitch labeling with other FLIM and PLIM assays (Fig. 5). In this experiment, we collected fluorescence signals from Matrigel in blue (donor) and green (acceptor) spectral channels for ColBD-Twitch protein and green fluorescence from Lgr5-GFP regions in organoids. We also expected that in addition to GFP (τ∼2.4 ns) FLIM method would enable measuring other green dyes and biosensors due to the possibility of separation in lifetime (‘time’) domain. Thus, we successfully imaged live organoid oxygenation in such setup, using blue-excited red-emitting phosphorescent O_2_ probe, which has lifetimes in range of 20-57 μs [65] (Fig. 5A). With another probe, TMRM, labeling mitochondria and displaying orange emission and membrane polarization-dependence of fluorescence lifetimes [67], we were able to gather information on GFP, TMRM-FLIM and ColBD-Twitch-labeled Matrigel scaffold (Fig. 5B) and observed heterogeneous mitochondrial polarization in stem cell niche in agreement with previous findings [67].

**Figure 5.**
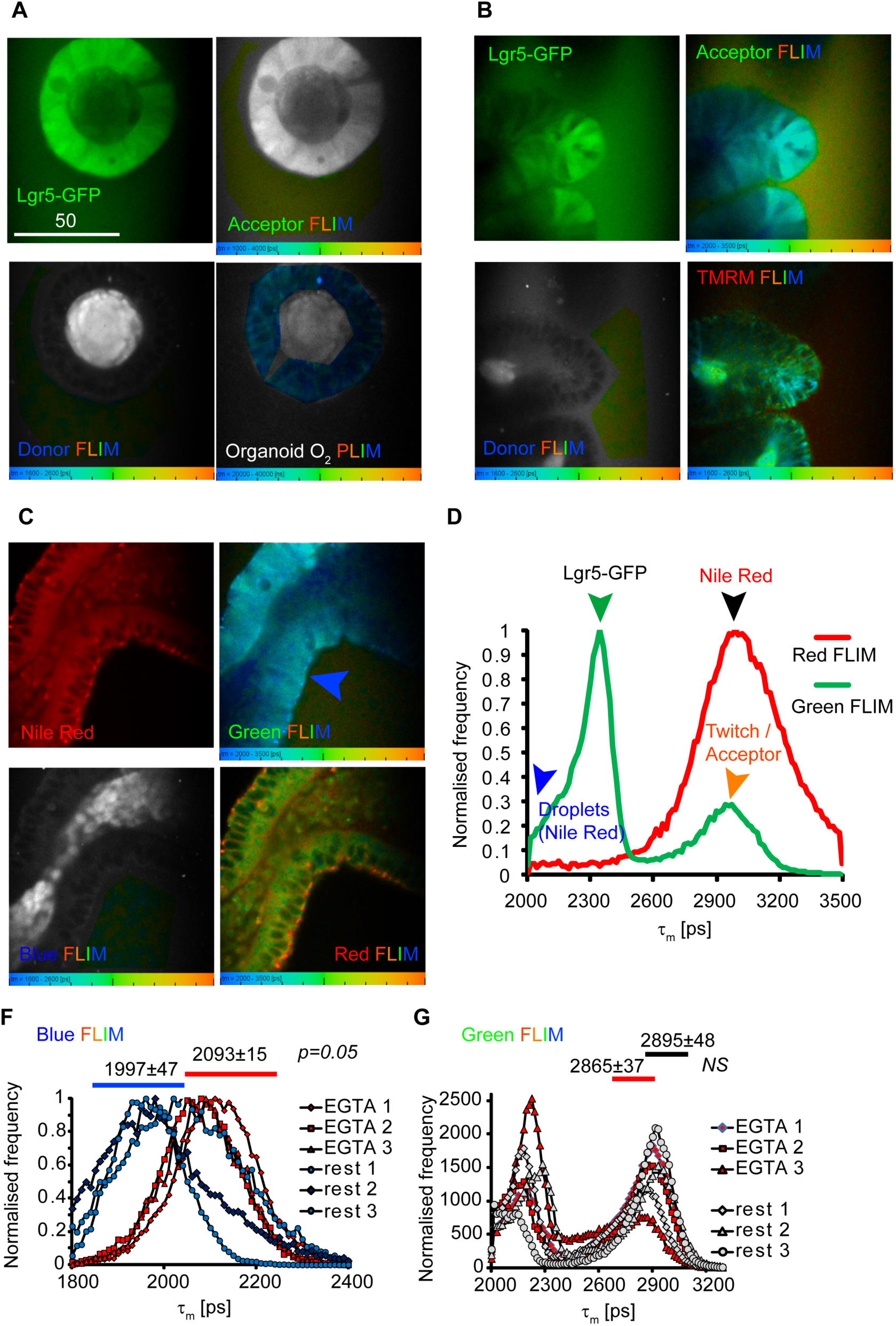
Examples of multi-parameter FLIM imaging of Matrigel stained with ColBD-Twitch in the culture of mouse intestinal Lgr5-GFP organoids. Organoids were grown in ENR VC medium and were stained with O_2_ probe (Pt-Glc, 2 μM, 1 h), TMRM (10 nM, 1 h) or Nile Red (1 μg/ml, 1 h). Matrigel was stained prior to measurements with ColBD-Twitch (2 μM, 1 h) in complete growth medium. A: Donor (Blue, 470 nm em.) and Acceptor (Green, 530 nm em.) FLIM images showing stained Matrigel, Lgr5-GFP (green, intensity) and O_2_-PLIM (635-675 nm em.). B: Lgr5-GFP, green and blue FLIM channels and TMRM-FLIM (488 nm exc., 565-605 nm em.). C, D: Nile Red (intensity, 565-605 nm em.), Green, blue and red FLIM images (C) and lifetime distribution histograms for respective green (512-536 nm) and red (565-605 nm) FLIM channels for Nile Red-stained organoid samples (D). Characteristic peaks of lifetimes for droplets in green channel (basal membrane), GFP, Matrix are indicated by arrows. F, G: comparisons of blue and green FLIM distribution histograms for VD3-stimulated (100 nM, 3 d) organoids in response to Ca^2+^ depletion (2.5 mM EGTA). Significant differences were observed only in blue (donor) channel. N=3. Scale bar is in μm.

We then chose more challenging probe, Nile Red, which is frequently used for assessment of lipids and lipid droplets; note that it also shows displays Ca^2+^-sensitivity, e.g. by detecting conformational changes of calmodulin upon binding Ca^2+^ [68, 69] or by binding phospholipid / Ca^2+^ complexes [70-72]. In the presence of lipids and other amphiphilic molecules, Nile Red displays both spectral (green / orange) and fluorescence lifetime changes. We tested if the green fluorescence of Nile Red could potentially interfere with Lgr5-GFP signals (Fig. 5C, D). While the fluorescence intensity signals in green channel (512-536 nm em., exc. 488 nm) overlapped, the lifetimes were distinguishable on a frequency lifetime distribution histogram (Fig. 5D green), showing three areas of lifetime distribution: <2.1 ns for basal membrane-localized droplets, ∼2.3-2.4 ns for Lgr5-GFP and ∼3 ns peak for the Matrigel labeled with ColBD-Twitch acceptor fluorescence. In contrast, Nile Red fluorescence in red channel (565-605 nm, ‘red FLIM’) demonstrated single peak of overlapping lifetimes in a range of 2.6-3.5 ns corresponding to membrane, cytoplasmic (lower lifetimes in 2.6-3 ns range) and lipid droplet (generally higher lifetimes in 2.8-3.5 ns range) staining (Fig. 5C and 5D red).

In a proof-of-principle experiment we tested responses of donor (blue) and acceptor (green) fluorescent forms of Matrigel-bound ColBD-Twitch to the depletion of the extracellular Ca^2+^ upon chelation with 2.5 mM EGTA. In agreement with calibration experiments (Fig. 3-4), the donor channel demonstrated most striking response to Ca^2+^ depletion (Fig. 5F,G).

Thus, staining of organoids with Nile Red allowed identifying fraction of basal membrane-localized lipid droplets, seen in both green and red spectral channels (Fig. 5, S6). Pre-treatment with 1α, 25(OH)_2_-vitamin D3 (VD3) affected both fluorescence lifetimes of Nile Red cytoplasmic and membrane-stained structures and increased the number of lipid droplets in organoids (Fig. S6). This effect supports previous report on the effect of VD3 on lipid metabolism in the intestine [73] and suggests that VD3 upregulates Ca^2+^ metabolism [74]. We then decided to investigate how the changes in extracellular Ca^2+^ could affect Nile Red fluorescence in the intestinal organoids.

We found that basal membrane-localized lipid droplets seen in different spectral channels did not fully co-localize and their fluorescence lifetimes tended to be different (Fig. 6A). We hypothesized that such droplets were of similar type, but the different lifetimes and spectral characteristics could indicate difference in labeling of their content by Nile Red. The short lifetimes in a green channel can reflect labeling of neutral lipid cores (mainly triacylglycerol and sterol esters) of lipid droplets, while the longer red fluorescence lifetimes can correspond both core and polar monolayers consisting of phospholipids and lipid droplet-associated proteins [68, 75]. We performed sequential addition of 0.5 and 2.5 mM EGTA and analyzed fluorescence lifetimes for Matrigel / ColBD-Twitch (blue channel) and for Nile Red (green and red channels) (Fig. 6B).

**Figure 6.**
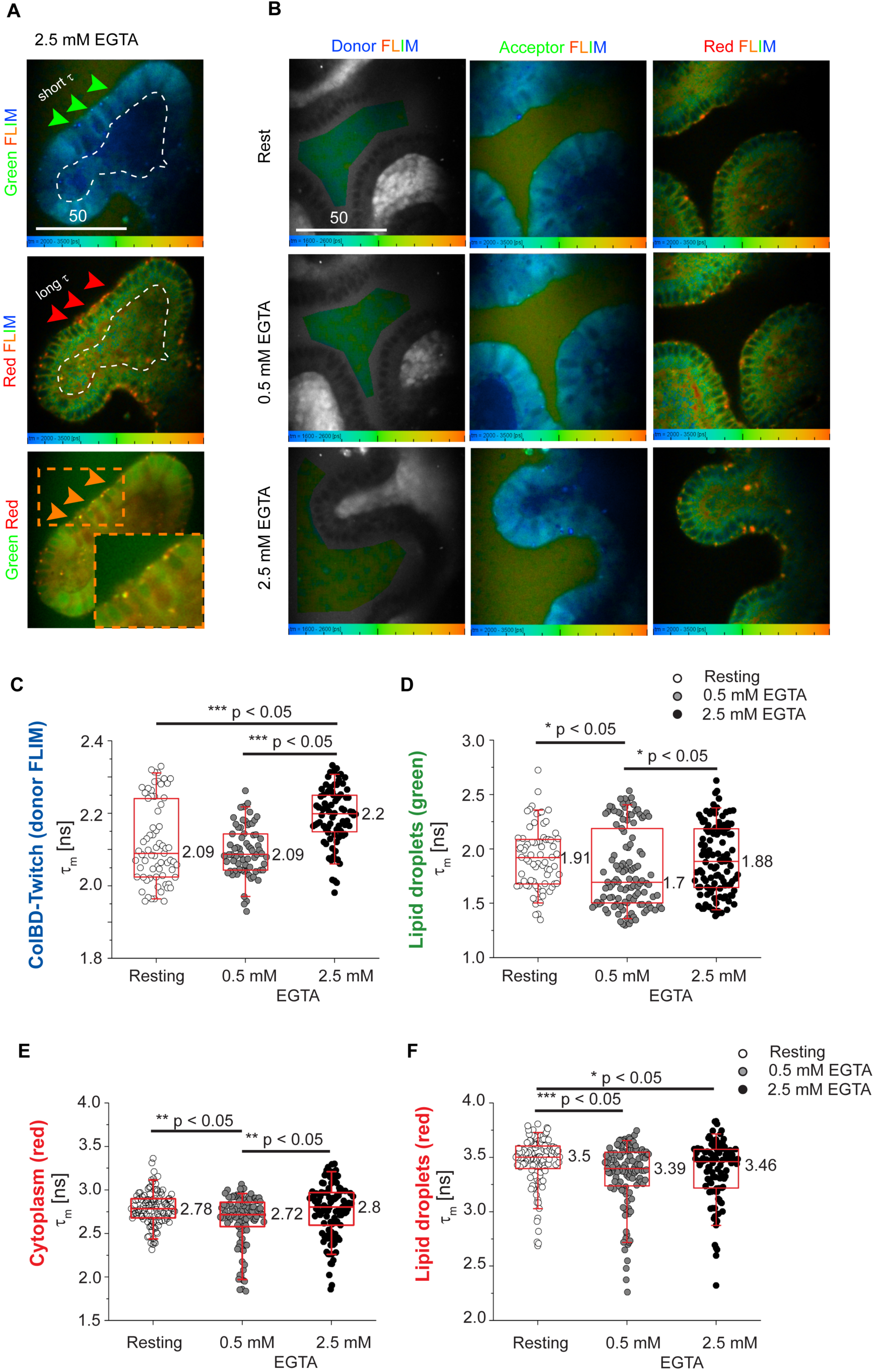
Multi-parameter FLIM of extracellular Ca^2+^ and lipid droplets in live mouse Lgr5-GFP intestinal organoids. A: basal membrane localization of Nile Red-stained lipid droplets in organoids. Green (512-536 nm em.) and red (565-605 nm em.) FLIM and combined intensity green/red images are shown. Droplets are indicated by arrows. Lumen is indicated by white dashed line. B: Examples of spectrally resolved (Donor, 446-486 nm em., Acceptor, 512-536 nm em. and red channel, 565-605 nm em.) FLIM images of Nile Red-stained Lgr5-GFP organoids and ColBD-Twitch-stained Matrigel matrix at rest and after sequential addition of EGTA (5 min). C-F: General statistical analysis of fluorescence lifetimes for donor FLIM channel (Matrix) measured with ColBD-Twitch, cytoplasmic (red), green and red spectral fractions of Nile Red measured and calculated at rest and different concentrations of EGTA (first experiment). The data points correspond to average lifetime values calculated from ROIs taken from 7 organoid images per each condition. P values indicate statistical significance (Mann-Whitney test): * - p<0.05, ** - p<0.005 and ***-p<0.0005. Box charts correspond to median (values shown in numbers), 25 and 75 percentiles. Whiskers show 5 and 95 percentiles. Scale bar is in μm.

First, we found that ColBD-Twitch did not detect statistically significant changes (Fig. 6C) or detected statistically decrease (Fig. S7A) rather than expected increase of donor fluorescence lifetime between resting (0 mM EGTA) and 0.5 mM EGTA concentrations. However, it worked successfully at 2.5 mM EGTA in both experimental replicates, demonstrating significant increase of a donor fluorescence lifetime (Fig. 6C and S7A).

Interestingly, analysis of Nile Red-stained droplets (green and red spectral channels, Fig. 6D,F) and membrane/cytoplasmic Nile Red signals (Fig. 6E) showed significant response to Ca^2+-^depletion already after addition of 0.5 mM EGTA. These results were not reproduced in the second experimental replicate (Fig. S7B, C), but showed the same tendencies. Thus, sequential chelation of external Ca^2+^ affected Nile Red fluorescence clearly demonstrating changes in the intracellular environment. This is likely due to the conformational reorganization of lipid membrane and protein structures the cytoplasm.

In order to address the variability in Nile Red fluorescence lifetime response and understand why ColBD-Twitch demonstrated an increase rather than decrease of free extracellular Ca^2+^ after addition of 0.5 mM EGTA to the organoids, we performed time-dependent analysis of the fluorescence lifetime time changes (Fig. 7). This enabled us to detect very minor changes in lifetime values, which were not detectable by the ‘general’ statistical analysis. Analysis of such profile for ColBD-Twitch demonstrated that with the first EGTA addition the initial fluorescence lifetimes were growing (Fig. 7A) reflecting decrease of the extracellular Ca^2+^ concentration. These changes were visible on a lifetime distribution histograms for most organoids in the group (Fig. 7B). Such response of ColBD-Twitch accompanied the increased number of lipid droplets with longer fluorescence lifetimes in red and green spectral channels (Fig.7C, D). In contrast, organoids measured 7-8 min later after the first addition of EGTA displayed statistically significant decrease of ColBD-Twitch fluorescence lifetimes accompanied by the corresponding changes of Nile Red lipid droplets fluorescence (red and green channel) (Fig. 7A, C, D). This ‘drop’ of fluorescence lifetimes explains why we detected almost no changes or statistical decrease of ColBD-Twitch and Nile Red lifetime values using the ‘general statistical analysis’ approach (Fig. 6C-F, S7). This phenomenon can be explained by the ability of cells in organoids to regulate extracellular Ca^2+^ via control of Ca^2+^ efflux. In this case, the concomitant changes in the Nile Red fluorescence lifetimes provide an indirect evidence for such regulation.

**Figure 7.**
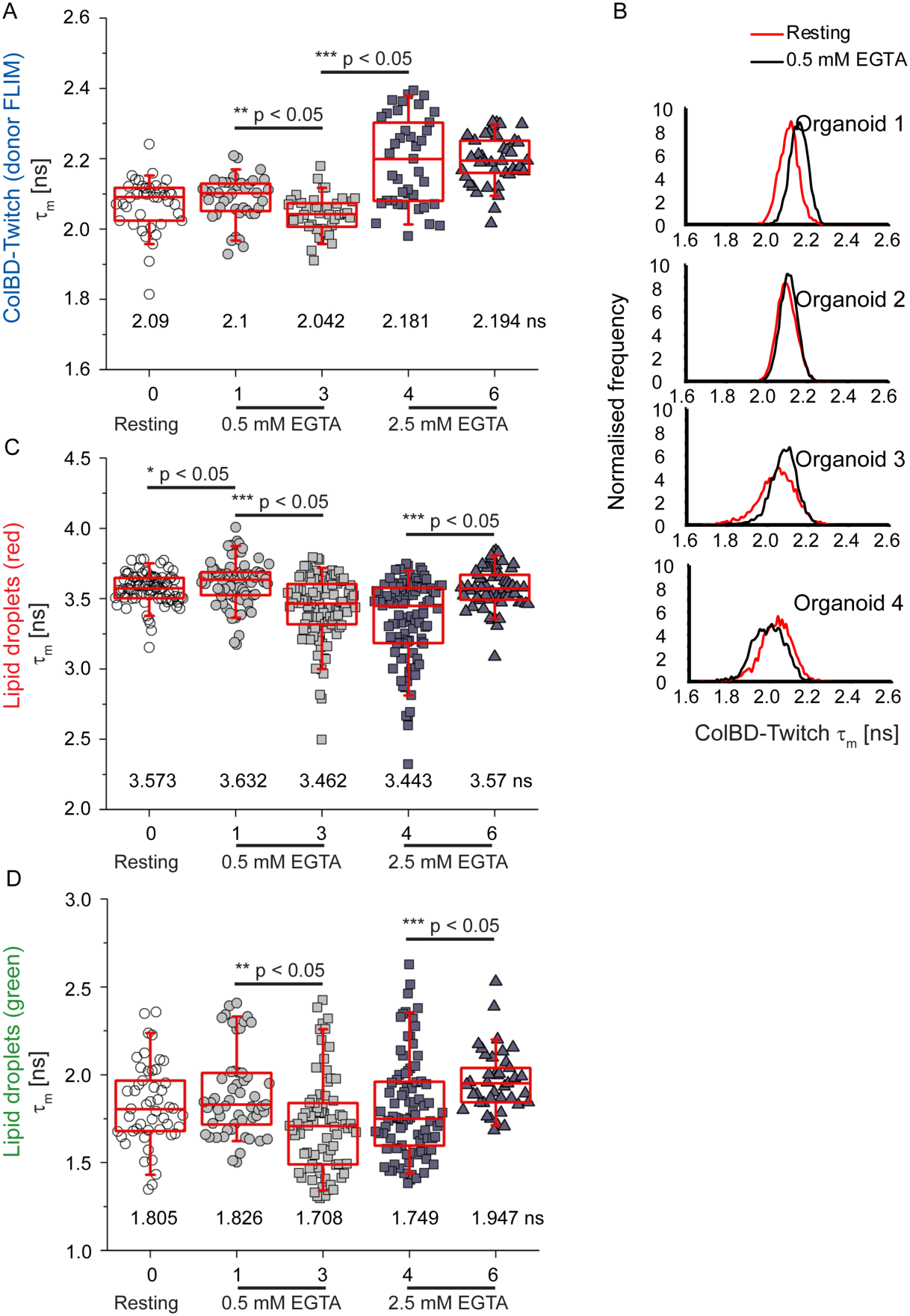
Time-dependent analysis of changes of fluorescence lifetimes for donor FLIM channel (ColBD-Twitch / Matrix, A) and different spectral fractions of Nile Red (C, D) in response to addition of EGTA. Data are combined from the two independent experiments and show average lifetime values calculated from ROIs taken from 4 organoid images per each condition. P values indicate statistical significance (Mann-Whitney test): * - p<0.05, ** - p<0.005 and ***-p<0.0005. Box charts correspond to median (values shown in numbers), 25 and 75 percentiles. Whiskers show 5 and 95 percentiles. **B:** fluorescence lifetime distribution histograms of individual organoids at rest and immediately after addition of 0.5 mM EGTA for different ColBD-Twitch-stained Matrigel areas.

Adding more EGTA led to the statistical increase of ColBD-Twitch fluorescence lifetime values due to decrease of free extracellular Ca^2+^ (Fig. 7A), which correlated with slow evolving increase of lipid droplet fluorescence lifetime in green and red spectral channels (Fig. 7C, D). Although we did not observe further increase of fluorescence lifetime after the second addition of EGTA with ColBD-Twitch, the cells could still attempt restoring balance of chelated extracellular Ca^2+^: the delayed response of Nile Red fluorescence could be viewed as evidence of activation of the intracellular Ca^2+^ pools in response to addition of 2.5 mM EGTA.

The involvement of organoid epithelium in regulation of extracellular Ca^2+^ was confirmed in the control experiment, where we treated organoid-free Matrigel with the sequential addition of EGTA (Fig. S8).

## Discussion

Here, we describe design and application of the extracellular Ca^2+^-sensing fluorescent protein, which provides labeling of the extracellular matrices for measurements in 3D tissue models and in related *in vivo* settings. While the sensing principle representing binding domain combined with fluorescent protein biosensor is not new, this is the first use of FRET-FLIM biosensor as a collagen-based scaffold biosensor and its successful application.

Recombinant cellulose- and collagen-binding domain-based Twitch-2B chimeras showed high expression yields (7-60 mg/L culture) and efficient folding when expressed in bacteria, especially in case of ColBD-Twitch protein fused to the short vWF peptide. Thus, we achieved improved production and demonstrated versatility of the approach suggesting that similarly other matrix-binding polypeptides can be employed for other types of scaffolds and related applications. Produced proteins showed very plausible scaffold binding properties and reminiscent to the previously described CBD-ECFP and cellulose [76].

We found that binding domain attached to Twitch-2B displayed minimal or no negative effects on the performance of biosensor in a ratiometric mode, which is highly similar to the biosensors reported in previous study [53]. On the other hand, Ca^2+^-sensitivity in a fluorescence lifetime domain has not been reported before for the Twitch proteins: as expected with similar types of FRET-FLIM biosensors [28, 52, 54], donor fluorescence decay displayed double-exponential decay, with Ca^2+^-dependent changes in A1 fraction and FRET efficiency [77]. Thus, Ca^2+^-calibration can be realized in units of mean lifetime τ_m_ (double-exponential fit), A1 [%], FRET efficiency and by other means. In our case, we found τ_m_ as the most useful readout reflecting robust and reproducible changes in response to Ca^2+^, under one- and two-photon excitation modes (Fig. 3, 4). On the other hand, acceptor fluorescence lifetime did not show reliable changes in response to Ca^2+^ (Fig. 4, 5). This can be explained by less efficient energy transfer from Cerulean to cpVenus due to differences in the protein microenvironment such as pH leading to dissipation of the energy or the stability of cpVenus as acceptor. Indirectly, this data points at FLIM as a preferred readout for quantitative measurements with Twitch-2B 54S+ rather than the ratiometric intensity. This can be different from the situation with the endogenously expressed biosensor when its stability is not so strongly dependent on the environment and it is continuously synthesized and ‘refreshed’ by the cell [53].

Using of vWF peptide for labeling collagen scaffolds also suggests that the biosensor binding to the Matrigel and related matrices can potentially compete with Ca^2+^-sensing osteonectin / SPARC protein [78], which is known to be present in Engelbreth-Holm-Swarm tumor [79]. Therefore, it would be very interesting to test other collagen-binding domains for targeting Twitch or related biosensors to understand their binding and biosensing properties when bound to the Matrigel matrix, especially their influence by the Ca^2+^-dependent conformational changes of the scaffold.

Sensing in a fluorescence lifetime domain provides more options for spectral and lifetime-based multiplexing, and we demonstrated compatibility of Ca^2+^-FLIM by ColBD-Twitch with assays measuring cell oxygenation (O_2_-PLIM), mitochondrial polarization (TMRM-FLIM) and analysis of lipids and lipid droplets (Nile Red, based FLIM) using stem cell-derived Lgr5-GFP mouse small intestinal organoid model (Fig. 5-6).

Generally, sensing extracellular dynamic micro-gradients of Ca^2+^ using *in vitro* models is expected to have limited application potential. However, in our model we saw measurable changes of Ca^2+^ in the solution surrounding the scaffold in decellularized collagen-coated cellulose and in Matrigel matrices. We did not detect subtle differences in ‘Ca^2+^ gradients’ surrounding the organoids due to the non-perfect optical sectioning and rather slow and equilibrium-based measurements occurring within minutes. However, we showed usefulness of the sensor for combined measurements of Ca^2+^-dependent changes of lipid droplets with Nile Red. By using ColBD-Twitch as a reference marker of Ca^2+^ depletion, we observed dynamic changes in lipid droplets organization potentially linked to the local polarity changes of lipid structures detected by Nile Red (Fig. 6). These changes were correlated with extracellular Ca^2+^ fluctuations showing that extracellular and intracellular Ca^2+^ pools are in close connection with each other. It should be kept in mind that use of Nile Red does not discriminate between fluctuations of free Ca^2+^ and Ca^2+^ bound to the phospholipids. As Ca^2+^-signaling regulates many different cellular functions in a wide temporal and concentration ranges, the intracellular fluctuations of free Ca^2+^ are strictly regulated and sequestered in the organelles or extracellular environment [21]. Potentially, observed changes in lipid droplet and cytoplasmic fractions of Nile Red were due the result of Ca^2+^ binding to the lipid droplets and cellular membranes, which could act as additional depots in cells of the intestinal epithelium, similar to cardiomyocytes, neuroepithelial and immune cells [71, 72, 80]. If this is the case, lipid droplets can play a role in rapid restoration and maintenance of the external Ca^2+^ pool.

Our data provides basis for future study of processes of lipid and Ca^2+^ metabolism in the intestinal organoids; logical next step would be a more detailed analysis of droplets and Ca^2+^ dynamics by comparing kinetical (< 1 s) and equilibrium responses. The performed extracellular Ca^2+^ analysis would benefit from parallel measurement of the intracellular Ca^2+^ in intestinal organoids, but we did not perform these measurements using Oregon Green BAPTA-1 [81] FLIM probe here, as we found that it did not stain Lgr5-GFP organoids embedded in Matrigel.

Importantly, observed performance of cellulose and collagen-binding domain Twitch-2B biosensors highlights their potential for advanced *in vivo* and *ex vivo* measurements, such as microinjection in live zebrafish or such mouse tissues as cartilage, bone marrow and intestine. When mixed with collagen or cellulose-based matrices, these biosensors can be also highly useful for *ex vivo* analysis of tumor organoids, wound healing studies or more complex stem cell-derived organoid models.

## Materials and methods

Antarctic phosphatase and restriction enzymes were purchased from New England Biolabs (Brennan & Co, Dublin, Ireland). T4 DNA Ligase, 2X polymerase chain reaction (PCR) buffer, Wizard Mini-Preps Plasmid DNA purification and SV Gel Clean-up kits were purchased from Promega (MyBio, Ireland). The O_2_-sensitive phosphorescent probe Pt-Glc was synthesized as described before [82]. DC celery scaffolds were prepared essentially as described previously [33].

Tetramethylrhodamine methyl ester (TMRM), Nile Red, CellLytic B reagent, protease inhibitors, lysozyme, Bradford protein assay reagent, LB Broth and all the other reagents were from Sigma-Aldrich (Dublin, Ireland). Standard sterile TC plates and plastic were from Sarstedt (Germany), Collagen I-coated plates and half-area plates were from Greiner (Cruinn, Dublin, Ireland). Microfluidic channel slides μ-Slide VI^0.1^ and multi-well silicone inserts were from Ibidi GmbH (Martinsried, Germany).

### Molecular cloning and production of recombinant proteins

Plasmid DNA encoding Twitch-2B 54S+ low affinity mutant generously provided by Prof. O. Griesbeck was prepared and described previously [53]. Insert encoding the ORF was PCR-amplified using primers 5’-GATCGGTACCGGTGGTGGTTCTGGTGGTATGGTGAGCAAGGGCGAGG-3’ (forward) and 5’-AGTCGGTACCTCAATCCTCAATGTTGTGACGGA-3’ (reverse) and cloned using *Kpn*I site into CBD Δ(FTN) vector essentially as described before [33], in order to produce CBD-Twitch expression construct. For ColBD-Twitch, we used custom-synthesized (Genscript, USA) sequence encoding N-terminal 6xHis tag, vWF peptide WREPSFMALS and Twitch-2B 54S+ (cloned using *Kpn*I site). All the produced expression constructs were verified using sequencing (GATC Biotech AG). Expression constructs were propagated in *E.coli* SG13009 cells, expressed (0.125 mM IPTG, room temperature, 18 h for ColBD-Twitch and 0.25 mM IPTG, 4 °C, 72 h for CBD-Twitch) and purified essentially as described previously with CBD-ECFP protein [76]. Purified proteins were dialyzed against PBS and stored at 4 °C. Total protein concentration was assessed using Bradford assay (Sigma) and spectrophotometrically (considering ε_515_ = 83,000 M^-1^ cm^-1^ for Ca^2+^-free form of cpVenus). Typical protein yields were 7 mg /L (CBD-Twitch, 49-80% folding rate) and >60 mg/ L (ColBD-Twitch, 55-70% folding rate).

### Scaffolds and coatings

Various types of collagen-based matrices were used in study: 96-well flat bottom plates or DC celery coated with collagen IV from human placenta diluted as described in [83], black-walled collagen I pre-coated 96-well plates (Greiner 655956) and Matrigel (‘growth factor reduced’ grade, Corning, 356231) distributed either as monolayer (15 μl per well of half-area plates, Greiner 675161) or drop-cast on 35 mm microscopy imaging dishes. Proteins (CBD-Twitch or ColBD-Twitch) were diluted either in phosphate-buffered saline (PBS, pH 7.4), 0.135 M NaCl or organoid growth media, added to the collagen or Matrigel matrices and incubated for 0.5-16 h at room temperature before analysis.

### Mouse intestinal organoid culture

Mouse Lgr5-GFP organoids were cultured as described previously [66]. To stain the Matrigel, ColBD-Twitch was added to the organoid culture in a complete growth medium (1 μM,1 h), washed once and imaged in Phenol Red-free DMEM medium (Sigma-Aldrich, D5030) supplemented with 10 mM HEPES, pH 7.2, 10 mM D-glucose, 2 mM L-glutamine and 1 mM sodium pyruvate. For co-staining and multiplexed FLIM-PLIM, we used following additional probes: Pt-Glc (2 μM, 1 h), TMRM (10 nM, 0.5 h), Nile Red (1 μg/ml, 1 h). For VD3 treatment, organoids were incubated with 100 nM for 3 days prior to measurements. For titration, EGTA was added sequentially at final concentrations 0.5 and 2.5 mM. The solution of EGTA (45 mM) was prepared from original 450 mM stock solution, pH 8.0 using tenfold dilution in the imaging media.

### FLIM microscopy

**Confocal FLIM** was performed on a custom-made upright Zeiss Axio Examiner Z1 upright laser-scanning FLIM-PLIM microscope as described previously [66, 76]. The microscope was equipped with 5x/0.25 Fluar air, 20x/1.0 and 63x/1.0 W-Plan Apochromat water-dipping objectives, integrated T and Z-axis controls, BDL-SMNI 405 and 488 nm pulsed diode lasers, DCS-120 confocal TCSPC scanner, photon counting detectors and SPCM and SPCImage software (Becker&Hickl GmbH). Lgr5-GFP, Pt-Glc (PLIM) and TMRM (FLIM) probes were imaged as described previously [66, 67], Nile Red was excited using 488 nm laser, emission collected at 512-536 nm (‘green’, Semrock) and 565-605 nm (‘red’) spectral channels. ColBD-Twitch was excited either with 405 nm (em. 446-486 nm, ‘Donor’ fluorescence, Semrock) or 488 nm (em. 512-536 nm, ‘Acceptor’ fluorescence).

FLIM calibration experiments were typically performed using Matrigel stained with ColBD-Twitch in PBS or 0.135 mM NaCl supplemented with different concentrations of CaCl_2_ and measured using 5x/0.25 Fluar air objective (405 nm and 488 nm excitation).

**Multiphoton FLIM** measurements were performed using an inverted Zeiss LSM 880 microscope (Zeiss, Jena, Germany) coupled with a Ti:Sapphire laser (Mai Tai HP; Spectra Physics, Santa Clara, USA) and a two-channel ‘BIG 2.0’ FLIM detector (Becker&Hickl GmbH). Images were collected with a 25x/0.8 NA objective (LCI Plan-Neofluar; Zeiss, Jena, Germany), 10% laser power and a pixel dwell time of 16.38 µs. Multiphoton excitation and emission curves were measured using the Lambda mode at a 8.5 nm channel resolution. FLIM measurements of DC celery scaffolds coated with collagen IV and stained with ColBD-Twitch were performed at 860 nm with a resolution of 256×256 pixels. The collected signal was separated using splitting into two channels, 460-500 nm for channel 1 and 520-560 for channel 2.

### Spectral and plate reader measurements

Absorption and fluorescence spectra of CBD-Twitch and ColBD-Twitch in solution were collected on 8453 diode array spectrophotometer (Agilent) and LS-50B (PerkinElmer) at room temperature as described previously [76]. Staining efficiency and ratiometric fluorescence measurements in solution and in collagen scaffolds were performed using BMG Clariostar microplate reader (BMG Labtech Ltd, UK) equipped with atmospheric control unit (CO_2_, T, O_2_), monochromator and MARS software.

### Data analysis and statistics

Fitting of FLIM decays was performed using SPCImage software, essentially as described previously, using mono- (phosphorescence lifetime imaging) and double-exponential decay fitting procedures (considering χ^2^<1.5 factor as appropriate). FRET efficiency and A1 (Figs. 3-4) were calculated as E [%] = 1-τ_1_*/*τ_2_ using SPCImage software.

Typically, fitting was tested for few separately taken pixels and applied to the whole 256×256 (or 512×512) image. Lifetime frequency distribution histograms were used either for chosen image regions (e.g. areas with Matrigel or basal membrane-localized lipid droplets) and used for further analysis. The peak values (identified using integration function in Origin software) were tested for normality of distribution (Shapiro-Wilk test) and evaluated for statistical significance of differences using *t*-test (normal distribution) or Mann-Whitney tests, where appropriate.

The titration of Matrigel / intestinal organoids with EGTA was performed twice, using cultures from different passages (Figs. 6-7, S7). Each of experimental replicate consisted of two independent treatments, thus four independent treatments were performed for EGTA experiment. For general statistical evaluation, we compared data collected from all microscopy images corresponding to each treatment and the measurements performed on different organoid passages were analyzed separately (Fig. 6C-F, S7). ROIs of the same size corresponding to the independent lipid droplets (for green or red spectral channels of Nile Red) or labeled Matrigel areas (blue spectral channel of ColBD-Twitch) collected from independent microscopy images were assumed as individual statistical units. The set of data for each experimental condition were analyzed for normality of distribution and with the non-parametric Mann-Whitney test. For a time-dependent analysis (Fig. 7), we combined microscopy images of organoids from four independent treatments with EGTA and organized them into 5 groups (by 4 organoids in each group) according to the order of imaging (Resting: organoids measured prior to the addition of 0.5 mM EGTA, 0.5 mM EGTA (1) and 2.5 mM EGTA (4): organoids measured immediately after addition of EGTA, 0.5 mM EGTA (3) and 2.5 mM EGTA (6): organoids measured 7-8 min after addition of EGTA) using ROIs collected from these images as statistical units as described above.

## Acknowledgments

This work was supported by the Science Foundation Ireland (SFI) grant 18/IF/6238 and Russian Science Foundation (RSF) grant 18-15-00407 (‘production of scaffolds’ part).

We are grateful to Prof. O. Griesbeck (Max Plank University) for providing Twitch-2B 54S+ expression construct and Dr. E. Grebenik (Sechenov University, Institute for Regenerative Medicine, Moscow) for providing Collagen-Chitosan matrix for the initial tests and to Prof. D. Papkovsky (University College Cork) for providing the access to FLIM microscope.

