## Supplementary Figures S1-S8 for "Extracellular Ca^2+^-sensitive fluorescent protein biosensor based on a collagen-binding domain"

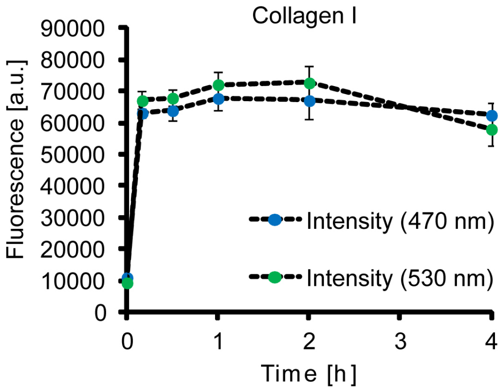


**Figure S1.** Staining of collagen with ColBD-Twitch protein. Collagen I-coated plates were incubated with 0.5 μM ColBD-Twitch, washed and measured on a microplate reader in two spectral channels (470 nm and 530 nm, as indicated).

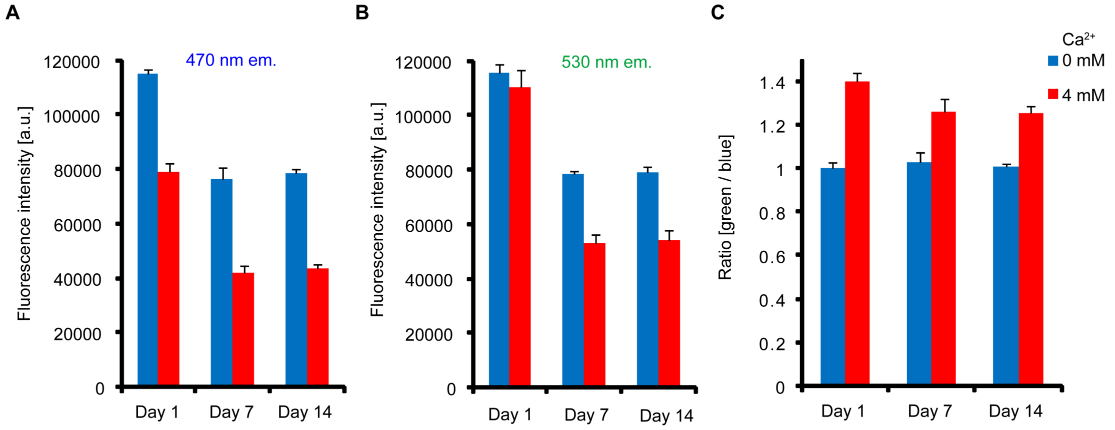


**Figure S2.** Storage stability of ColBD-Twitch protein. Sample was diluted to 0.1 μM, measured on a microplate reader and incubated at 4 °C before analysis at days 7 and 14 at 470 and 530 nm. Ratio (530/ 470) is also shown. N=3.

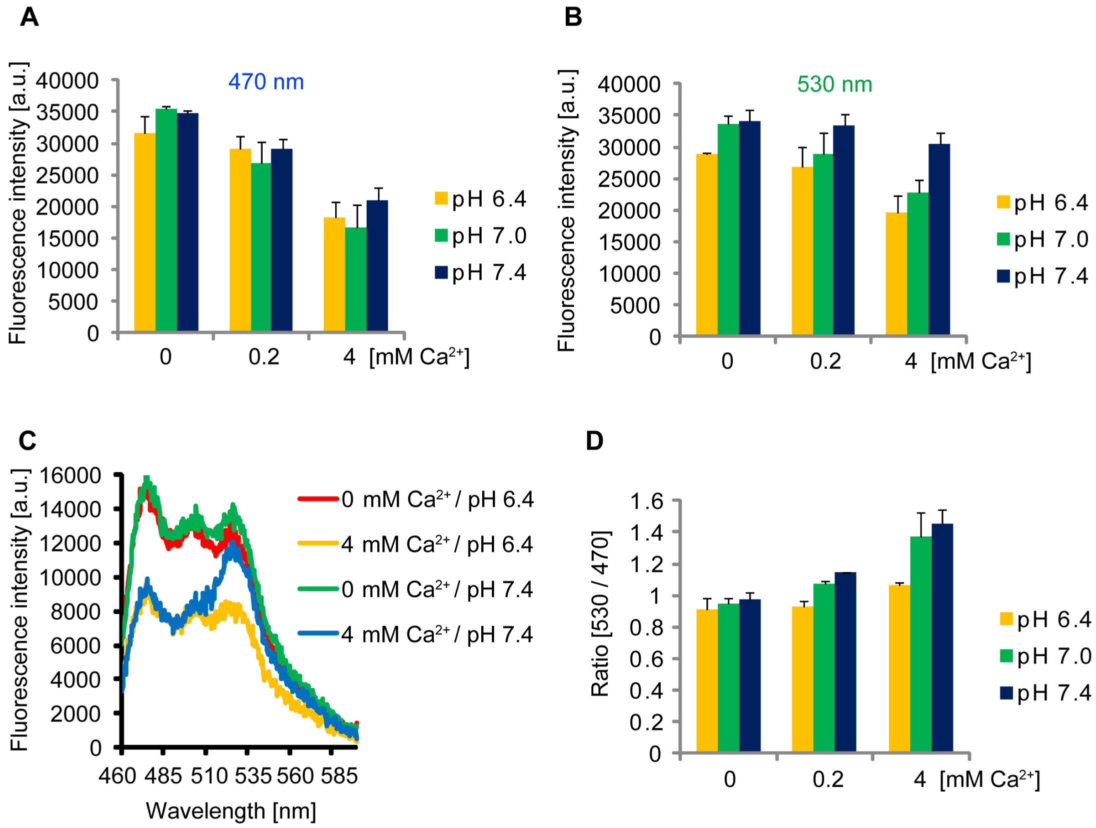


**Figure S3.** pH sensitivity of ColBD-Twitch protein. The protein was diluted to 0.07 μM in buffers with different pH and concentrations of $CaCl_2$ as indicated and measured on a microplate reader at 470 nm (A), 530 nm (B) and in spectral emission scan modes (C). D: pH-dependence of ratio response. N=3.

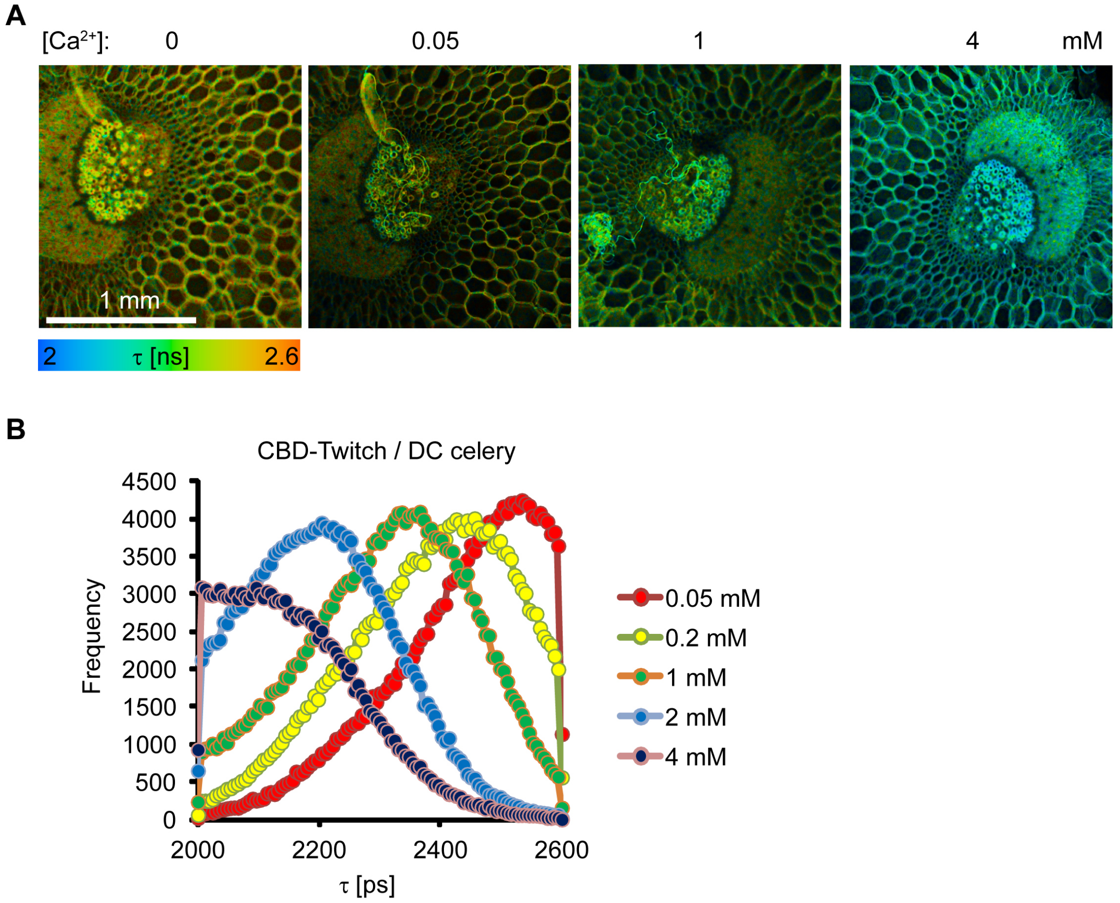


**Figure S4**. $Ca^{2+}$-dependent FLIM response of decellularized celery matrix stained with CBD-Twitch protein (1 μM in HBSS, overnight incubation) and measured in the presence of different concentrations of $CaCl_2$. Examples of FLIM images (A) and normalized lifetime frequency distribution histograms (B) are shown.

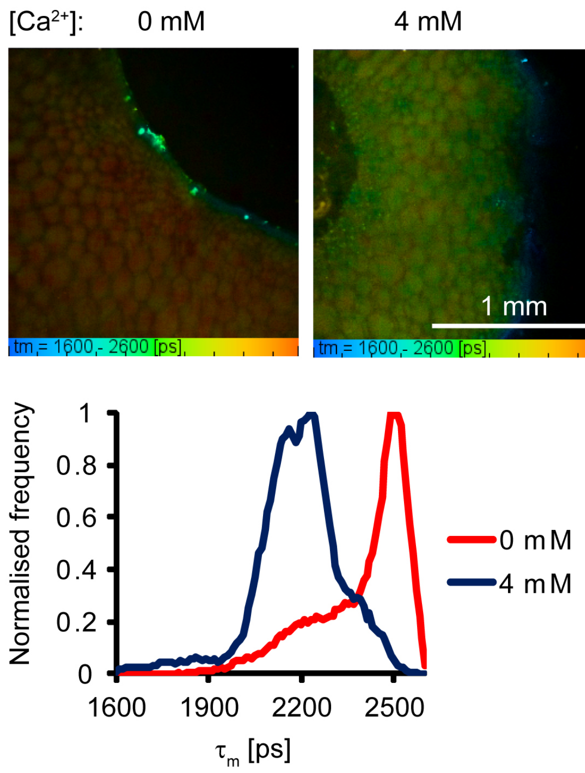


**Figure S5.** $Ca^{2+}$-dependent FLIM response with decellularized celery matrix pre-coated with Collagen IV (0.1%, overnight incubation) and stained with ColBD-Twitch protein (1 µM, overnight). Produced matrix was subsequently imaged on FLIM (405 nm exc., 468 nm em.) at 0 and 4 mM $CaCl_2$ in PBS. Below: normalized fluorescence lifetime distribution histograms.

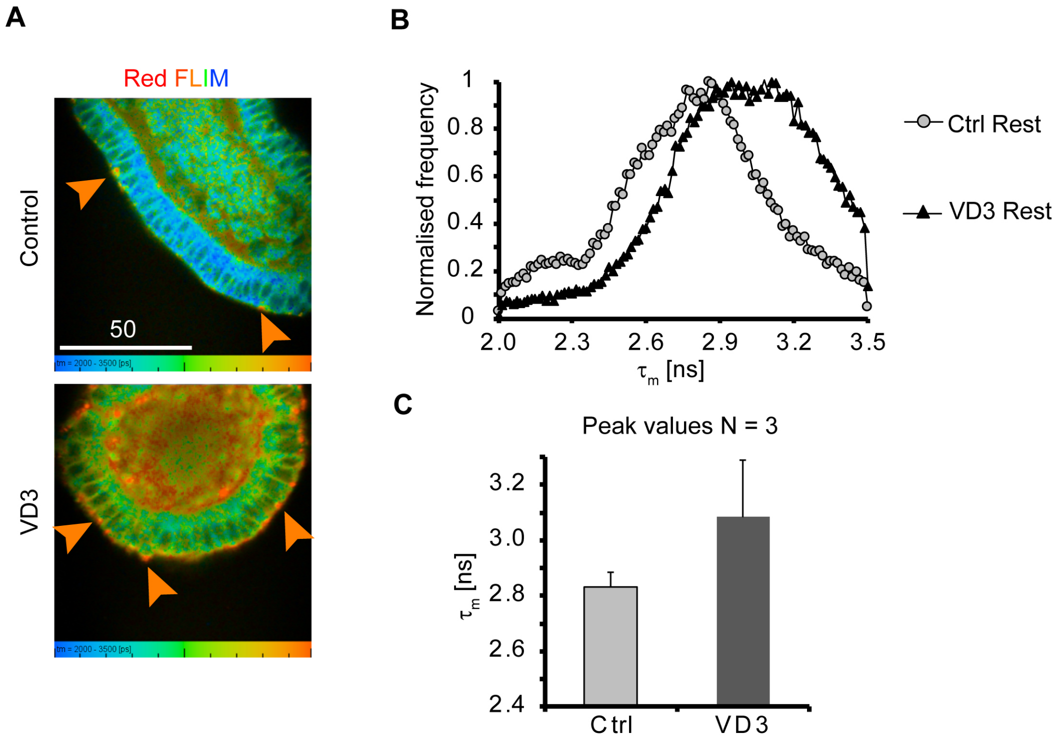


**Figure S6:** FLIM of Nile Red-stained intestinal organoids displays $Ca^{2+}$ and VD3-dependent responses. Organoids were grown in presence of VD3 (100 nM, 3 d) and treated with 2.5 mM EGTA. A: FLIM images in red fluorescence channel. B, C: Lifetime distribution histograms and comparison of peak values for different experimental conditions. Scale bar is in µm.

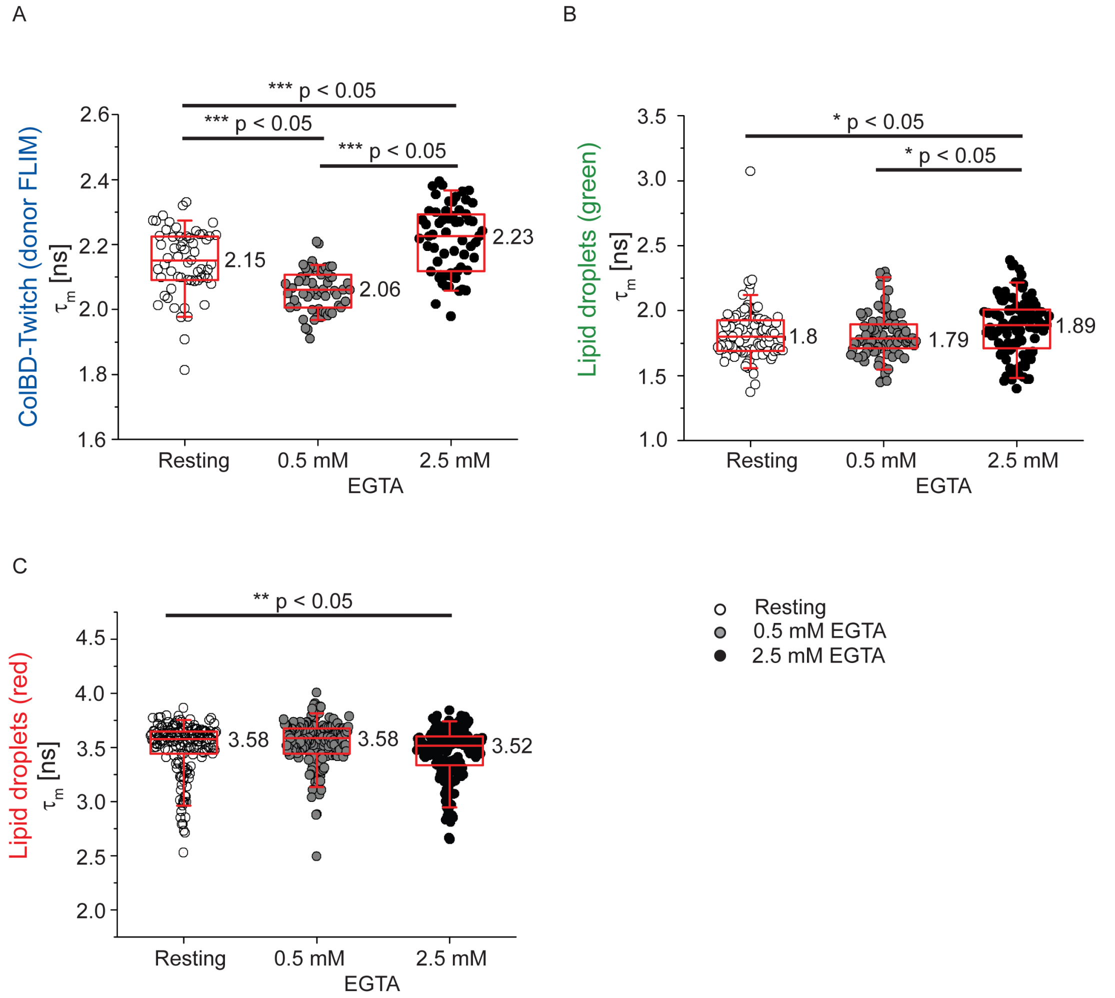


**Figure S7.** General statistical analysis of fluorescence lifetimes for donor FLIM channel (Matrix) measured with ColBD-Twitch (a), green (b) and red (c) spectral fractions of Nile Red measured and calculated at rest and different concentrations of EGTA added (**second experiment**). The data distributions correspond to the average lifetime values calculated from ROIs taken from 6 organoid images per each condition. P values indicate statistical significance (Mann-Whitney test): * - p<0.05, ** - p<0.005 and ***-p<0.0005. Box charts correspond to median (values shown in numbers), 25 and 75 percentiles. Whiskers show 5 and 95 percentiles.

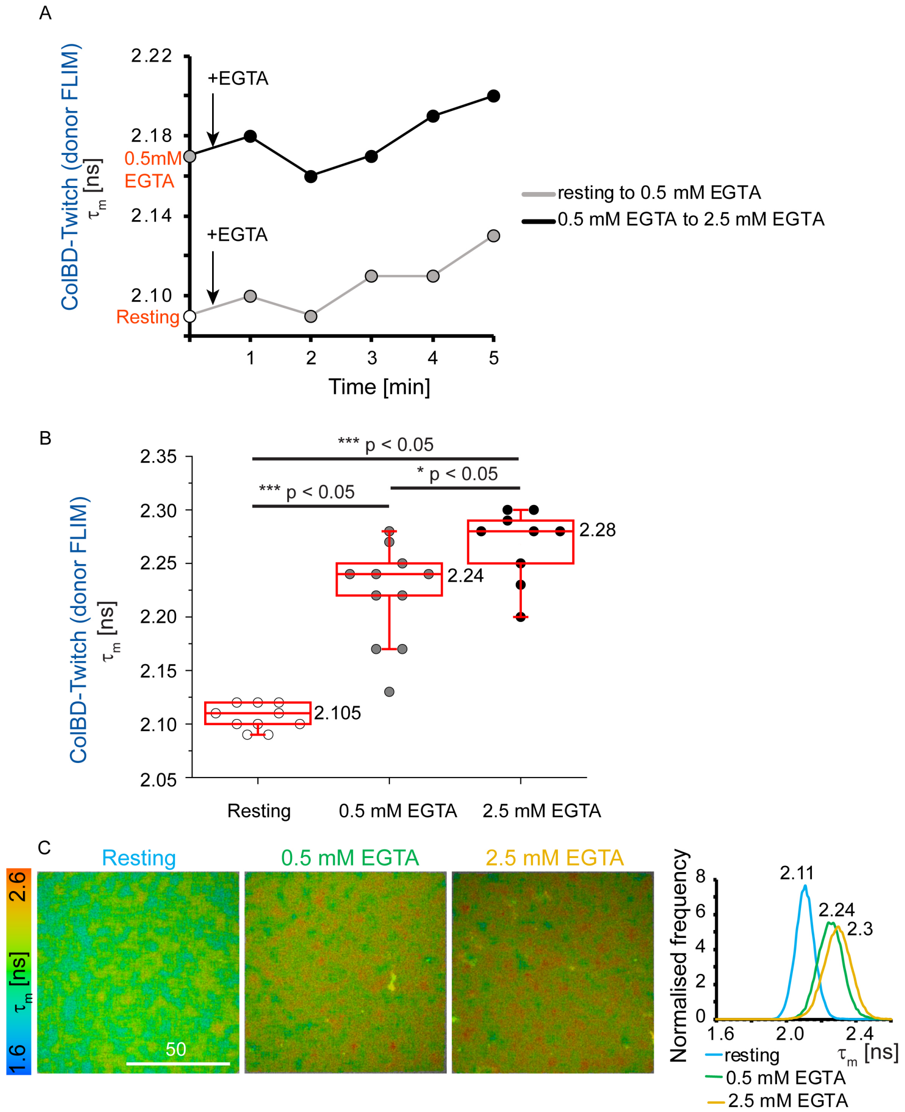


**Figure S8.** Changes in fluorescence lifetime for ColBD-Twitch bound to the Matrigel and treated with EGTA in the absence of organoids. A: Time-dependent changes of ColBD-Twitch (donor)-stained Matrigel fluorescence. The kinetical profiles are shown for two independent ROIs. Data corresponds to the peak values of distribution histograms measured for the same ROI prior to addition of EGTA (resting or 0.5 mM EGTA) and at different time points. B: General statistical analysis of fluorescence lifetimes at resting, in presence of 0.5 mM EGTA and 2.5 mM EGTA. Data correspond to the peak values of fluorescence lifetime distribution histograms for different ROIs measured at rest and after EGTA addition. P values indicate statistical significance (Mann-Whitney test): * - $p<0.05$, ** - $p<0.005$ and ***-$p<0.0005$. Box charts correspond to median (values shown in numbers), 25 and 75 percentiles. Whiskers show 5 and 95 percentiles. C: Examples of FLIM images of stained Matrigel and their corresponding distribution histograms. Scale bar is in μm.
